# Genomic miscellany and allelic frequencies of *Plasmodium falciparum msp-1, msp-2* and *glurp* in parasite isolates

**DOI:** 10.1101/2021.06.25.449861

**Authors:** Ibrar Ullah, Sahib Gul Afridi, Asif Ullah, Muhammad Israr, Anwar Ali, Hina Jabeen, Akhtar Rasool, Fazal Akbar, Muzafar Shah

## Abstract

The genomic miscellany of malaria parasites can inform the intensity of transmission and identify potential deficiencies in malaria control programs. The aim of this study was to investigate the genomic miscellany, allele frequencies and multiplicity of infection (MOI) of *P. falciparum*.

**Methods:** A total of 85 isolates from patients presenting to the local health centers with *P. falciparum* species were collected from 2017 to 2019. Parasite DNA was extracted from a total of 200 µl whole blood per patient using the Qiagen DNA extraction kit according to manufactures instructions. The polymorphic region of *msp-1, msp-2* and *glurp* loci were genotyped by using nested polymerase chain reactions followed by gel electrophoresis for fragment analysis.

**Results:** Genetic diversity and allelic frequencies of *msp-1, msp-2* and *glurp* were identified in 85 blood samples. A total of 62 *msp* alleles were detected in which 30 for *msp-1* and 32 for *msp-2*. For *msp-1* the successful amplification occurred in (75/85) 88.23% isolates for *msp-1*, 78.9% (67/85) for *msp-2* and 70% (60/85) for *glurp*. For *msp-1*, the K1 allelic family was predominant at 66.66% (50/75), followed by RO33 and MAD20. The frequency of samples having only K1, MAD20 and RO33 were 21.34% (16/75), 8% (6/75) and 10.67% (8/75) respectively. In *msp-2*, the FC27 allelic family was the most abundant with 70.14% (47/67) compared to 3D7 with 67.16% (45/67). Nine *glurp* RII region genotypes were identified. The overall mean multiplicity of infection was 2.6 with1.8 and 1.4 for *msp-1* and *msp-2* respectively while for *glurp* RII genes (MOI=1.03). There was no significant association between multiplicity of infection and age group (Spearman’s rank coefficient = 0.050; *P* = 0.6). There was significant correlation between MOI and parasite density for *msp-2* allelic family.

**Conclusion:** Our study showed high genetic diversity and allelic frequency with multiple clones of *msp-1, msp-2* and *glurp* in *P. falciparum* isolates from malaria patients in Khyber Pakhtunkhwa Pakistan. In the present study the genotype data provided the valuable information which is essential for monitoring the impact of malaria eradication efforts in this region.

## Background

Human malaria is a blood infectious disease caused by mosquito borne apicomplexan parasites of the genus *Plasmodium* and it is mediated by the arthropod vector *Anopheles* mosquito. This parasite is unicellular eukaryotes that invades host erythrocytes and reside within a parasitophorous vacuole (Bonnefoy *et al*., 2008). Among the five species of malaria parasite, *P. falciparum* is the fatal species show high mortality and exhibits complex genetic polymorphism which may be explain its ability to develop multiple drug resistance and avoid vaccines (Wongsrichanalai *et al*., 2002).

Pakistan is one of the malaria endemic countries in Asia for both *P. vivax* and *P. falciparum* malaria (Asif, 2008 & Yasinzai *et al*., 2008). Pakistan shares boarders with countries like Afghanistan, India and Iran, where the disease is also endemic. There is large cross boarder human migration within these regions which also increase refuges influxes, all of which facilitate malaria transmission. The introduction of new type of alleles by migration, mutation and recombination creates genetic diversity in population while on the other hand immune responses of the human host as well as chemotherapy play key roles in selection which affects the frequency of new alleles in parasite population (Ord *et al*., 2005).

Molecular epidemiological studies remain an important tool to analyze the genetic diversity of *P. falciparum* population especially in areas of intense malaria transmission. The strategy to control malarial infection requires an understanding of the genetic composition of *P. falciparum* as this information is essential and may facilitate the development of an effective anti-malarial vaccine (Olasehinde *et al*., 2012, Hamid *et al*., 2013 & Kiwuwa *et al*., 2013). In order to determine the number and the types of parasite clones, genotyping is an important tool to determine malaria parasite populations. In molecular epidemiological studies of malaria, this approach is used to analyze the genetic diversity of infections with consideration of different factors like transmission intensity and host immunity. The most widely used techniques for malaria genotyping are based on amplification by PCR of the polymorphic genes encoding the merozoite surface proteins *msp-1, msp-2* and the glutamate rich protein (*glurp*) (Snounou *et al*., 1999 & Bakhi *et al*., 2015). The *msp-1* and *msp-2* are two main *P. falciparum* blood stage malaria vaccine targets (Chitarra *et al*., 1999) which show a very essential role in identification of genetically distinct *P. falciparum* parasite populations. The *msp-1* is a 190 KDa surface protein encoded by *msp-1* genes located on chromosome number 9 and contains 17 blocks of sequences flanked by conserved regions (Takala *et al*., 2002 & Tanabe *et al*., 1987). In *msp-1* (block 2) which is highly polymorphic part of *msp-1* is grouped into three allelic families namely K1, MAD20 and RO33 (Farrar *et al*., 2015). The *msp-2* is also polymorphic glycoprotein encoded by the *msp-2* genes located on chromosomes 2 and composed by five blocks (Smythe *et al*., 1991). The *msp-2* alleles are grouped into two allelic families FC27 and 3D7 (Mohammed *et al*., 2015). The *glurp* is an exo-antigen of *P. falciaprum* on which Phase I vaccine trials have been completed and it is expressed in both the pre-erythrocytic and erythrocytic stages of the parasite life cycle (Borre *et al*., 1991).

To our knowledge the genetic diversity of *P. falciparum* has been extensively studied in different parts of the world but there is very limited information on genetic diversity of *msp1, msp2* and *glurp* genes in our study site. This study aimed to evaluate the genetic diversity, multiplicity of infection, the level of malaria transmission and allele frequencies of *msp-1, msp-2* and *glurp* in malaria parasite isolated from different districts of our study area.

## Methods

### Study Site

This study was carried out in different districts of Khyber Pakhtunkhwa province. The latitude and altitude of study site is 34.9526 N and 72.3311 E. The population of study area is 2,203,000 (35.53 million) and the total area is 101741 km^2^. It receives the annual rain fall is 384 mm occurring in two seasons from March to May and from August to November. The mean temperature ranges from 20°C to 40°C. In the present study the annual parasite index (API) of KPK is 6.3 (WHO, 2020). In this region, *P. vivax* is the dominant malaria species. A sample of this study was provided by clinical trial conducted from 2017 to 2019 in different health facilities of nine different districts.

### Study population and blood sample collection

A total of 100 blood samples were collected from the patient having uncomplicated *P. falciparum*. Among 100 samples, 85 were positive for *P. falciaprum* by microscopy. The patient aged between 4 months to 60 years were resident of different district of the study site had presented to the local health center (Hospitals, clinics, laboratories, etc)with fever (≥37.5). Before blood samples collection, informed consent was obtained from all participants or in case of children from their parent guardians prior to their enrollment. This study was approved by BOS (Board of study) and ASRB (Advanced Studies Research Board) of Abdul Wali khan University Mardan. Venous blood (5ml) samples were collected in an EDTA (BD, USA) tube for microscopy examination and DNA extraction. The blood was stored at -20 C prior to DNA extraction at Biochemistry research lab, Abdul Wali Khan University Mardan, Pakistan. Genomic DNA was extracted from a total of 200 µl whole blood per patient using the Qiagen DNA extraction kit according to manufactures instructions. Nested polymerase chain reaction (PCR) genotyping was performed both for the variable block-2 regions of *msp-1, msp-2* and *glurp* genetic markers to assess potential diversity and allelic frequency of *P. falciparum* infection (Farnert *et al*., 2001).

### Allelic genotyping of *msp-1, msp-2* and *glurp* genes of *Plasmodium falciparum*

Nested PCR of the polymorphic region of *msp-1* (block-2), *msp-2* (block-3) and *glurp* (RII repeat regions) was performed using primers and methods as previously described (Snounou *et al*., 2002 & Ntoumi *et al*., 2000). The initial amplification primers pairs corresponding to conserved sequences within polymorphic regions of each gene were included in separate reactions. The product generated in the initial amplification was used as a template in separate nested PCR reactions. In the nested reaction separate primers pairs targeted the respective allelic types of *msp-1* (KI, MAD20 and RO33), *msp-2* (3D7 and FC27) and the RII block of *glurp* with amplification mixture containing 250 nM of each primer (except *glurp*nest 1 primers where 125 nM were used) 2mM of MgCl_2_ and 125 µM of each dNTPs and 0.4 units Taq DNA polymerase. The cyclic condition in the thermo cycler (My Cycler-Bio-Rad, Hercules, USA) for initial *msp-1* and *msp-2* PCR and initial nested *glurp* PCR were as follows: 5 min at 95C followed by 30 cycles for 1 min at 94°C, 2 min at 58°C and 2 min at 72°C and final extension of 10 min at 72°C. For *msp-1* and *msp-2* nested PCR conditions were follows: 5 min at 9°C, followed by 30 cycles for 1 min at 95°C, 2 min at 61°C and 2 min at 72 °C and final extension of 5 min at 72°C (Gosi P *et al*., 2013). The *msp1, msp2* and *glurp* fragments were amplified then resolved by gel electrophoresis in 2% agarose gel visualized under ultraviolet trans-illumination with light after staining with ethidium bromide. The size of DNA fragments were estimated by visual inspection using a 100 bp DNA ladder marker (New England Biolabs. Inc, UK)

### Malaria parasite identification

Thick and thin smears were stained with Giemsa. Parasite density was determined by analyzing the number of asexual parasites per 200 white blood cells and calculated per micro liter using the following formula which is the number of parasites x 8000/200 assuming the blood cell count of 8000 cells per µl. When there is no result on 200 high power ocular fields of the thick film, it was considered negative (WHO, 2015). The data were graded according to parasitaemia: 1, patients with 50-5000 parasites/µl; 2 patients with 5000 - 10000 parasites /µl; 3 patients with more than 10000 parasites /µl. (Mustafa *et al*., 2017) % parasitemia = Parasitized RBCs /total number of RBCs X 100

### Multiplicity of infection (MOI)

The mean multiplicity of infection (MOI) was calculated by dividing the total number of fragments detecting in both *msp-1* and *msp-2* loci by the number of sample positive for the both markers. The isolates with more than one allelic family were considered as polyclonal infection and those contain single allelic family were considered mono infection. The number of infecting genotypes contained in each sample which represents the multiplicity of infection (MOI) for that sample was calculated as a highest number of alleles obtained in the samples (Soe *et al*., 2017 & Duah *et al*., 2016).

### Statistical analysis

All the statistical analysis was performed by using the software SPSS version. The *msp1, msp2* and *glurp* allelic frequency was calculated. The mean MOI was calculated for *msp1, msp2* and *glurp* genes. The proportions of allele comparisons were assessed by Chi square tests. The spearman’s rank correlation coefficient was calculated to evaluate relationship between multiplicity of infection (MOI), parasite densities or age groups in the patients. A *P* value of ≤0.05 was considered indicative of statistically significant differences.

## Result

### General characteristics of study population

Overall, 100 randomly selected subjects of *P. falciparum* were screened, out of which 85 subjects were included in the clinical study. Out of 85 *P. falciparum* patients attending health facilities in study sites, 10.6% (09/85) patients were from Mardan, 12.94% (11/85) were from Swat, 11.8% (10/85) were from Buner, 11.8% (10/85) were from Hungo, 10.6% (9/85) were from Swabi, 11.8% (10/85) were from Kohat, 9.42% (08/85) were from Bannu, 10.6% (09/85) were from Timergara and 10.6% (09/85) were from Peshawar.The numbers of male patients were 58.8% (50/85) which were higherthan female patients 41.2% (35/85). The samples were collected from age 4 months to 65 years with a mean age of (33.9 ± 1.6) years. Among the entire participant in the age groups, the highest numbers of participant were noted in the age group (21-40) years which were 40% (34/85) and the lowest numbers of participants were noted in the age group (>60) years which were 1.2% (1/85). The mean body auxiliary temperature determined prior to blood sampling was (≥37°C). The parasite density ranged of 3451 to 89,045 parasites /µl with a mean density of 15197 parasites (95% CI (12059-18335) per µl of blood.The mean parasite density was higher in the Swat region (22303.45 Parasite/µl) as compared to Bannu which shows the least number of parasite densities (7623.37 Parasite/µl) (Table 1&2)

**Table 1:**
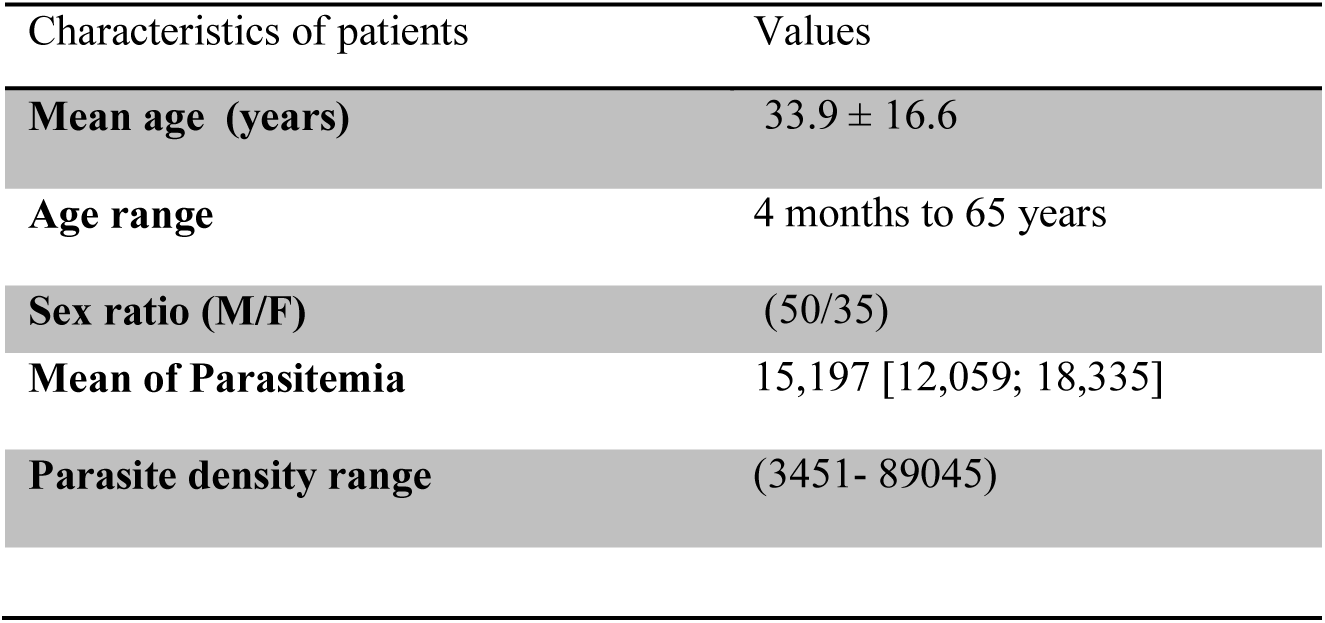
Demographic, data of the study population at districts, KP, Pakistan.

**Table 2:** Population characteristics of *P. falciparum* infected patients in the study area.

| Gender | Mardan<br>n (%) | Swat<br>n (%) | Buner<br>n (%) | Hungo<br>n (%) | Swabi<br>n (%) | Kohat<br>n (%) | Bannu<br>n (%) | Timergara<br>n (%) | Pesh<br>n (%) | Total |
| --- | --- | --- | --- | --- | --- | --- | --- | --- | --- | --- |
| **Male** | 7(77.7%) | 5(45.5%) | 4(40.0%) | 6 (60.0%) | 4(44.4%) | 6(60.0%) | 4(50.0%) | 7(77.8%) | 7(77.8%) | 50(58.8%) |
| **Female** | 2(22.3%) | 6(54.5%) | 6(60.0%) | 4 (40.0%) | 5(55.6%) | 4(40.0%) | 4(50.0%) | 2 (22.2%) | 2(22.2%) | 35(41.2%) |
| **Total** | 9(100.0%) | 11(100.0%) | 10(100.0%) | 10(100.0%) | 9(100%) | 10(100.0%) | 8(100.0%) | 9(100.0%) | 9 (100.0%) | 85(100.0%) |
| Age group |  |  |  |  |  |  |  |  |  |  |
| **<5** | 1<br>(11.11%) | 0 (0.00%) | 1 (10.0%) | 1 (10.0%) | 0 (0.00%) | 1(10.0%) | 1 (12.5 %) | 1 (11.11%) | 1 (11.11%) | 07 (8.24%) |
| **5-20** | 1<br>(11.11%) | 2 (18.18%) | 1 (10.0%) | 3 (30.0%) | 2 (22.22%) | 0 (0.0%) | 0 (0.00%) | 2 (22.22%) | 3 (33.33%) | 14 (16.47%) |
| **21-40** | 3<br>(33.33%) | 7 (63.63%) | 3 (30.0%) | 3 (30.0%) | 5 (55.55%) | 3(30.0%) | 2 (25.0%) | 4(44.44%) | 4 (44.44%) | 34 (40.0%) |
| **41-60** | 4<br>(44.44%) | 2 (18.18%) | 5 (50.0%) | 3(30.0%) | 2 (22.22%) | 5 (50.0%) | 5 (62.5%) | 2 (22.22%) | 1 (11.11%) | 29 (34.11%) |
| **>60** | 0 (0.00%) | 0 (0.00%) | 0(0.00%) | 0 (0.00%) | 0 (0.00%) | 1 (10.0%) | 0 (00.0%) | 0 (0.00%) | 0 (0.00%) | 01(1.18%) |
| **Total** | 9<br>(10.59%) | 11(12.95%) | 10 (11.76%) | 10 (11.76%) | 9 (10.59%) | 10(11.76%) | 8 (9.41%) | 9 (10.59%) | 9 (10.59%) | 85 (100.0%) |
| **Mean Parasitaemia** | 20233.83 | 22303.45 | 13639.8 | 9881.1 | 17358.5 | 15148.9 | 7623.37 | 19119.6 | 9856.3 |  |

### Frequencies of *msp-1, msp-2* and *glurp* allelic families

Out of 85 *P. falciparum* positive isolates, the successful amplification occurred in (75/85) 88.23% isolates for *msp-1*, 78.9% (67/85) for *msp-2* and 70% (60/85) for *glurp*. For *msp-1*, the K1 allelic family was predominant at 66.66% (50/75), followed by RO33 allelic family at 58.66% (44/75) and finally the MAD20 allelic family at 54.0% (41/75). Frequencies of different type of *msp-1* and *msp-2* alleles their combination and multiplicity of infection across the study sites are shown in (Table 3). The frequency of samples having only K1, MAD20 and RO33 were 21.3% (16/75), 8% (6/75) and 10.7% (8/75) respectively. Forty percent (40%) samples positive for *msp-1* were classified as monoclonal infection and the remaining sixty percent (60%) were classified as polyclonal infection with K1/RO33, K1/MAD20, and MAD20/RO33 representing 13.3% 12.00% and 14.7% respectively while 20.00% samples show all the three allelic type (trimorphic) together K1/MAD20/RO33. In *msp-2*, the FC27 allelic family was the most abundant with 70.14% (47/67) compared to 3D7 with 67.16% (45/67). The frequency for *msp-2*, the total monoclonal infections were (42/67) 62.68% in which the samples having only 3D7 allelic family were (20/67) 29.85% while the sample containing FC27 allelic type were (22/67) 32.8%. The samples with polyclonal infection were (FC27+ 3D7) was (25/67) 37.4%. For *glurp* 60 samples (70.6%) were found to be positive for RII repeats region producing 09 distinct sizes ranging between 600-1100bp among which 700bp allele was found to be predominant 21/60 (35%) (Table 3)

**Table 3:**
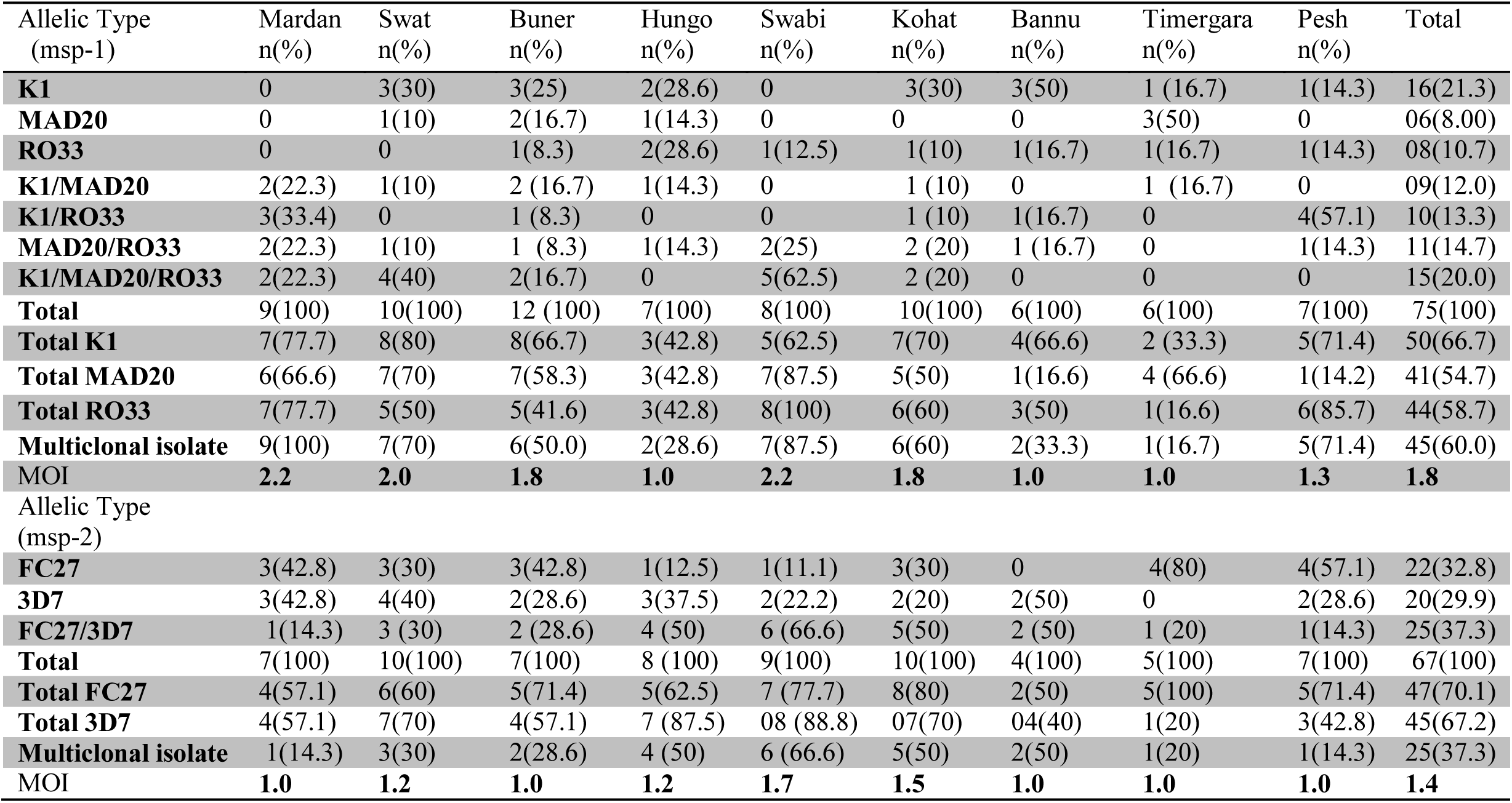
Frequencies and multiplicity of infection of Plasmodium falciparum isolates merozoite surface proteins-1 (*msp-1*), merozoite surface proteins-2 (*msp-2*) allelic forms.

| Allelic Type<br>(*msp-1*) | Mardan<br>n(%) | Swat<br>n(%) | Buner<br>n(%) | Hungo<br>n(%) | Swabi<br>n(%) | Kohat<br>n(%) | Bannu<br>n(%) | Timergara<br>n(%) | Pesh<br>n(%) | Total |
| --- | --- | --- | --- | --- | --- | --- | --- | --- | --- | --- |
| **K1** | 0 | 3(30) | 3(25) | 2(28.6) | 0 | 3(30) | 3(50) | 1 (16.7) | 1(14.3) | 16(21.3) |
| **MAD20** | 0 | 1(10) | 2(16.7) | 1(14.3) | 0 | 0 | 0 | 3(50) | 0 | 06(8.00) |
| **RO33** | 0 | 0 | 1(8.3) | 2(28.6) | 1(12.5) | 1(10) | 1(16.7) | 1(16.7) | 1(14.3) | 08(10.7) |
| **K1/MAD20** | 2(22.3) | 1(10) | 2 (16.7) | 1(14.3) | 0 | 1 (10) | 0 | 1 (16.7) | 0 | 09(12.0) |
| **K1/RO33** | 3(33.4) | 0 | 1 (8.3) | 0 | 0 | 1 (10) | 1(16.7) | 0 | 4(57.1) | 10(13.3) |
| **MAD20/RO33** | 2(22.3) | 1(10) | 1 (8.3) | 1(14.3) | 2(25) | 2 (20) | 1 (16.7) | 0 | 1(14.3) | 11(14.7) |
| **K1/MAD20/RO33** | 2(22.3) | 4(40) | 2(16.7) | 0 | 5(62.5) | 2 (20) | 0 | 0 | 0 | 15(20.0) |
| **Total** | 9(100) | 10(100) | 12 (100) | 7(100) | 8(100) | 10(100) | 6(100) | 6(100) | 7(100) | 75(100) |
| **Total K1** | 7(77.7) | 8(80) | 8(66.7) | 3(42.8) | 5(62.5) | 7(70) | 4(66.6) | 2 (33.3) | 5(71.4) | 50(66.7) |
| **Total MAD20** | 6(66.6) | 7(70) | 7(58.3) | 3(42.8) | 7(87.5) | 5(50) | 1(16.6) | 4 (66.6) | 1(14.2) | 41(54.7) |
| **Total RO33** | 7(77.7) | 5(50) | 5(41.6) | 3(42.8) | 8(100) | 6(60) | 3(50) | 1(16.6) | 6(85.7) | 44(58.7) |
| **Multiclonal isolate** | 9(100) | 7(70) | 6(50.0) | 2(28.6) | 7(87.5) | 6(60) | 2(33.3) | 1(16.7) | 5(71.4) | 45(60.0) |
| **MOI** | **2.2** | **2.0** | **1.8** | **1.0** | **2.2** | **1.8** | **1.0** | **1.0** | **1.3** | **1.8** |
| Allelic Type<br>(*msp-2*) |  |  |  |  |  |  |  |  |  |  |
| **FC27** | 3(42.8) | 3(30) | 3(42.8) | 1(12.5) | 1(11.1) | 3(30) | 0 | 4(80) | 4(57.1) | 22(32.8) |
| **3D7** | 3(42.8) | 4(40) | 2(28.6) | 3(37.5) | 2(22.2) | 2(20) | 2(50) | 0 | 2(28.6) | 20(29.9) |
| **FC27/3D7** | 1(14.3) | 3 (30) | 2 (28.6) | 4 (50) | 6 (66.6) | 5(50) | 2 (50) | 1 (20) | 1(14.3) | 25(37.3) |
| **Total** | 7(100) | 10(100) | 7(100) | 8 (100) | 9(100) | 10(100) | 4(100) | 5(100) | 7(100) | 67(100) |
| **Total FC27** | 4(57.1) | 6(60) | 5(71.4) | 5(62.5) | 7 (77.7) | 8(80) | 2(50) | 5(100) | 5(71.4) | 47(70.1) |
| **Total 3D7** | 4(57.1) | 7(70) | 4(57.1) | 7 (87.5) | 08 (88.8) | 07(70) | 04(40) | 1(20) | 3(42.8) | 45(67.2) |
| **Multiclonal isolate** | 1(14.3) | 3(30) | 2(28.6) | 4 (50) | 6 (66.6) | 5(50) | 2(50) | 1(20) | 1(14.3) | 25(37.3) |
| **MOI** | **1.0** | **1.2** | **1.0** | **1.2** | **1.7** | **1.5** | **1.0** | **1.0** | **1.0** | **1.4** |

### Genetic diversity and allelic frequency of *msp-1, msp-2* and *glurp* genes

The allelic genotyping display and show the polymorphic nature of *P. falciparum* in province of Khyber Pakhtunkhwa. In *msp-1, msp-2* and *glurp* different allelic types were identified. Alleles of *msp-1, msp-2* and *glurp* were classified according to the size of the amplified PCR bands. Samples amplification occurred in (75/85) 88.23% isolates for *msp-1*, 78.9% (67/85) for *msp-2* and 70% (60/85) for *glurp*. A total of 62 *msp* alleles were analyzed (30 for *msp-1* and 32 for *msp-2*) and 9 distinct alleles for *glurp* RII region. Thirty different alleles were noted in *msp-1*, among which 10 alleles for K1 (fragment range 100-400 bp), 11 alleles for MAD20 (fragment range 100-390bp) and 9 for RO33 (fragment range 100-360bp). In *msp-1*, K1 110 bp, 150 bp, 120 bp, MAD20 140 bp, 130 bp, 160 bp; and the RO33 120 bp, 130 bp, 140 bp were the most dominant (Figure 1, 4 & 5).

**Figure 1.**
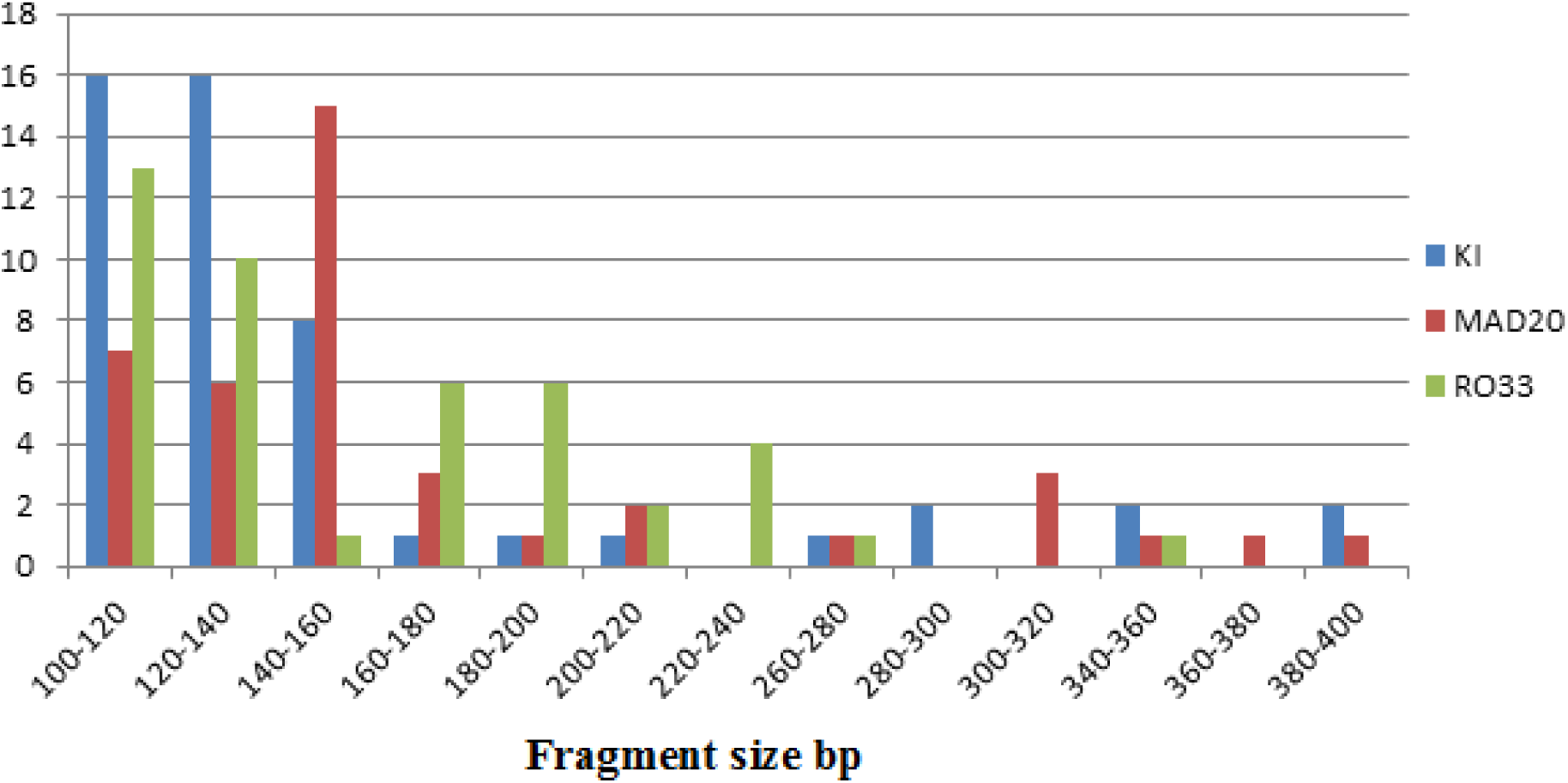
Prevalence of *P. falciparum* KI,MAD20, R033 *mspl* alleles classified by length

For *msp-2*, 32 individual alleles were noted with 16 alleles for FC27 (Fragment range 150-900 bp) in which 150 bp were predominant and 16 alleles for 3D7 (Fragment ranges 150-900bp) in which 150 bp were predominant (Figure 2).

**Figure 2.**
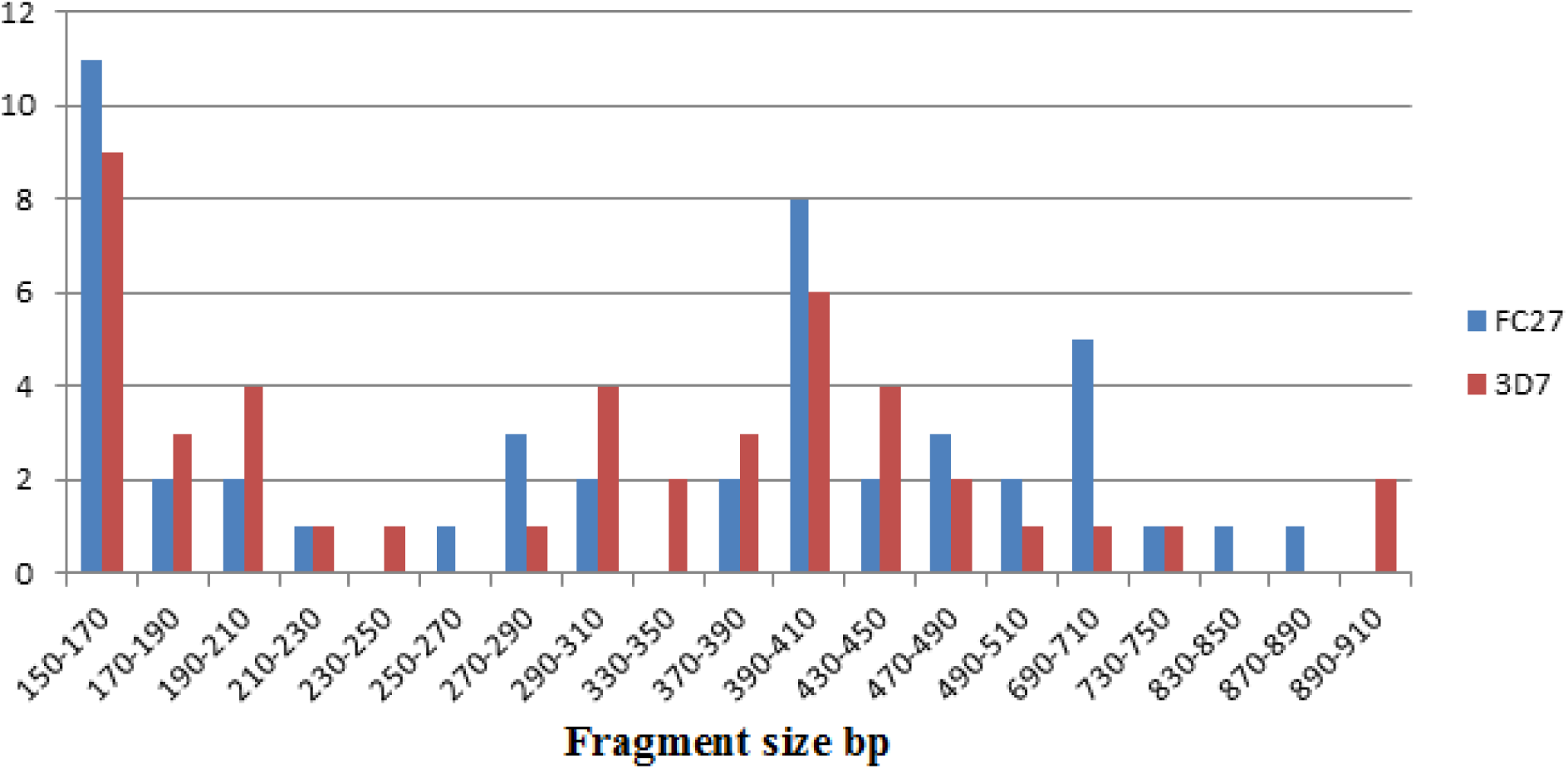
Prevalence of *P. falciparum* FC27 and 3D7 *msp2* alleles class.ified by length (base pair)

For *glurp*, 60 samples were genotyped for RII repeat region which produce 09 different alleles having size ranging from 600 to 1100 bp. These 9 alleles were coded as genotypes (1-9) among which genotype 2 (700 bp) 21/60 (35%) allele were found to be predominant, followed by genotype 3 and 8 (750 bp) (15%) and (1000 bp) (15%) (Table 4 & Figure 3)

**Table 4:** Distributions of allelic variants of *glurp* RII repeat region of *P. falciparum* populations in Khyber Pakhtunkhwa, Pakistan.

| Genotype | Allelic size variants (50bp bin ) | N (%) |
| --- | --- | --- |
| 1 | 600 – 650 | 07(11.7) |
| 2 | 651 – 700 | 21(35.0) |
| 3 | 701 – 750 | 09 (15.0) |
| 4 | 751 – 800 | 04( 6.6) |
| 5 | 801 – 850 | 03 (5.0) |
| 6 | 851 – 900 | 04( 6.6) |
| 7 | 901 – 950 | 01(1.6) |
| 8 | 951– 1000 | 09(15.0) |
| 9 | 1051- 1100 | 02 (3.33) |

**Figure 3.**
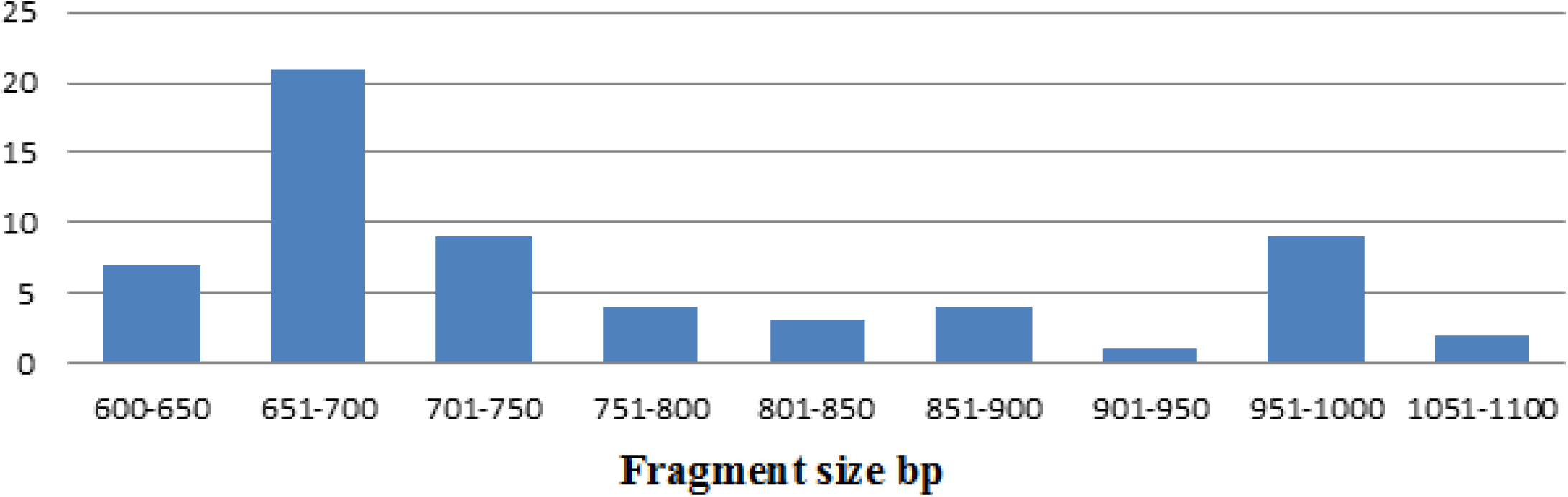
Prevalence of *P. falciparum glurp* alleles classified by length (base pair)

**Figure 4.**
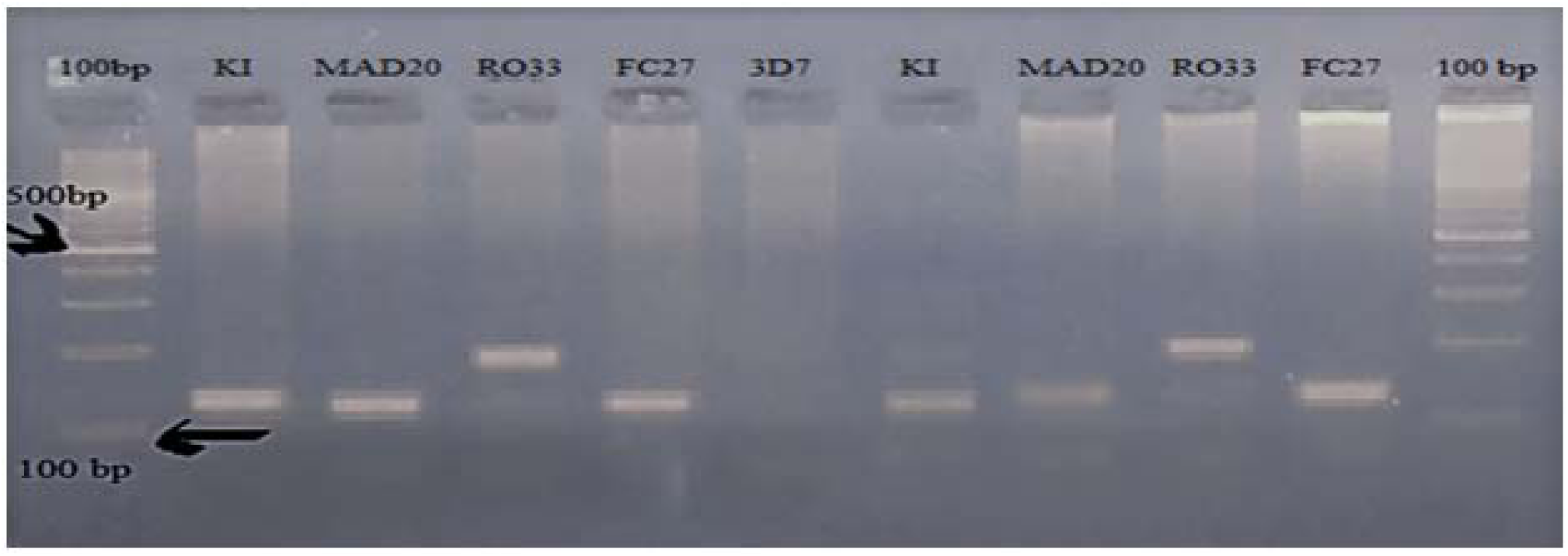
Gel picture showing the amplification of *P. falciparummsp-1* and *msp-2* genes. Lane 1 shows 100 bp DNA ladder. Lane 2 show’s *msp-1* allelic family KL lane 3 allelic family MAD20, lane 4 allelic family RO33 while for *msp-2*. Lane 5 and 6 show’s FC27 and 3D7 allelic family respectively.

**Figure 5.**
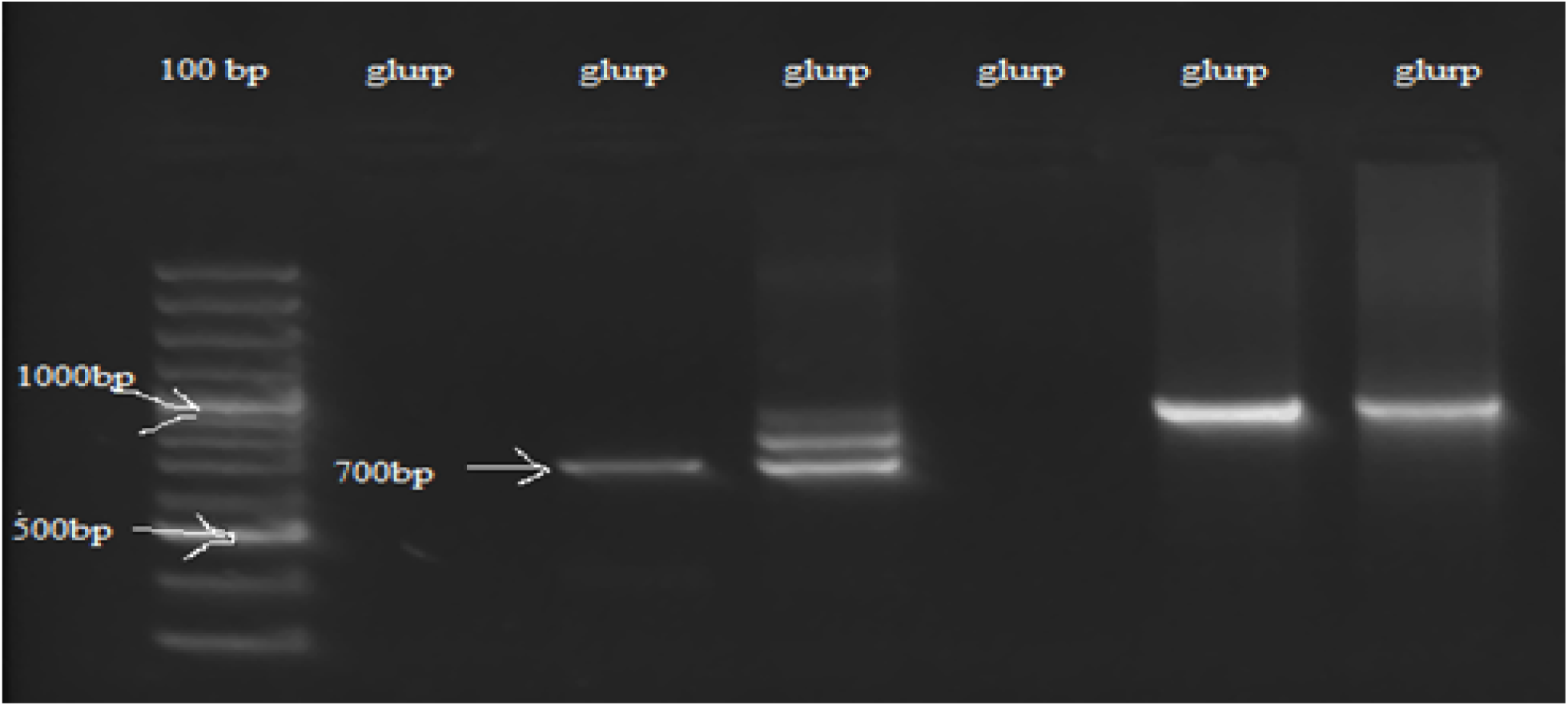
Agarose gel electrophoresis analysis showing genetic diversity detected with the *P. falciparum glurp* gene RII region. The *glurp* allele ranged in size from 600 bp-1100 bp. Marker: 100 bp ladder.

**Figure 6.**
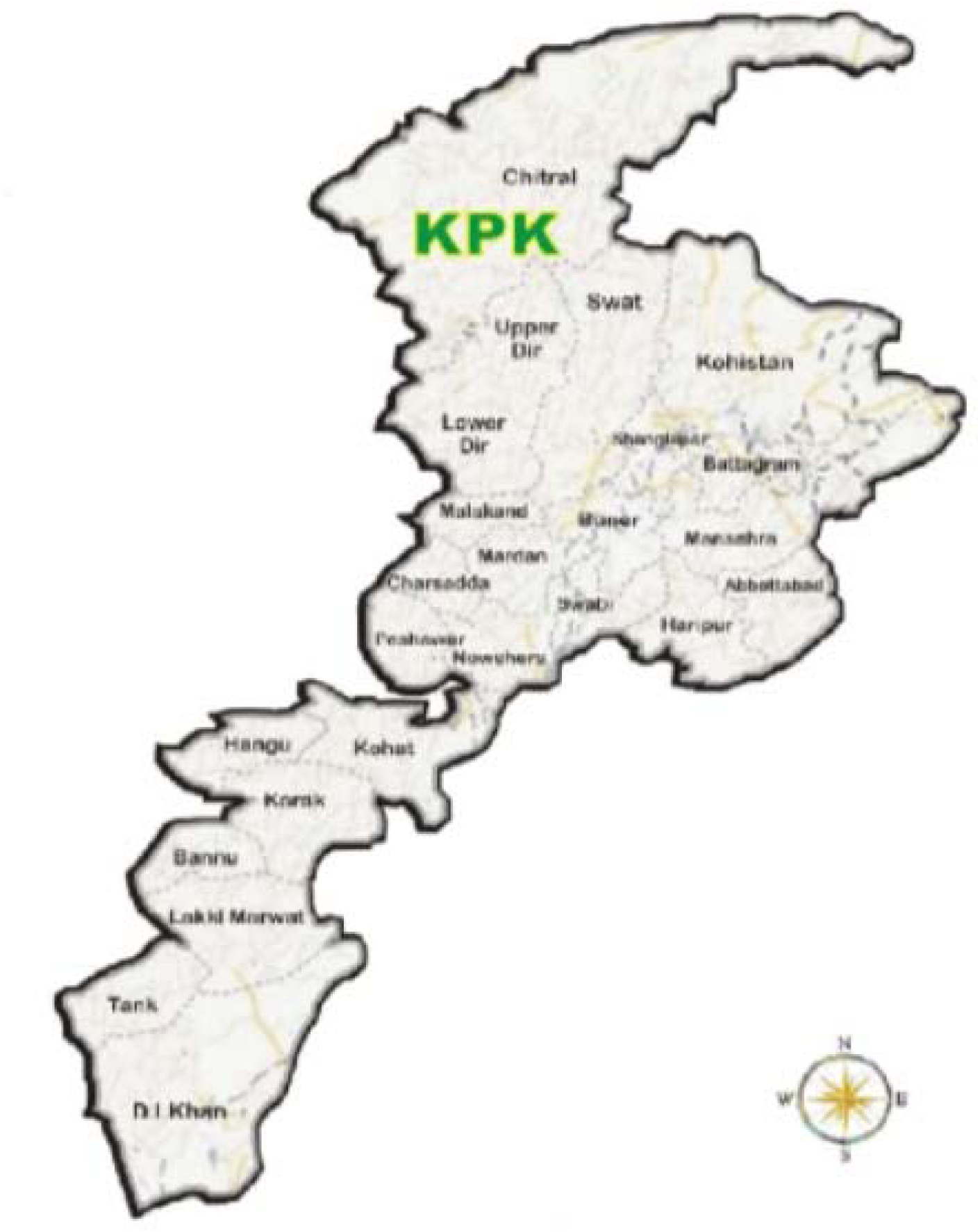
A geographic map showing different districts of Khyber Pakhtunkhwa Pakistan.

### Multiclonal Infection

The multiplicity of infection (MOI)/ number of genotypes per infection was calculated by dividing the total number of fragments detected in one antigenic marker by the number of samples positive for the same marker. Out of 85 samples (86.1%) show more than one parasite genotype. The overall mean multiplicity of infection was 2.6. Total of 60% (45/75) for *msp-1*, 37.3% (25/67) for *msp-2* and 3.34% (2/60) for *glurp* genes show multiple clones.The MOIs for *msp1, msp2* and *glurp* was 1.8 and 1.4 and 1.03 respectively.

### Relationship between multiplicity of infection, age groups and parasite densities

The prevalence of *msp-1, msp-2* and *glurp* allelic families and their combinations was significantly higher in number among patient age group 21-40 as compared to other age groups. However, there is no significant correlation between multiplicity of infection and age groups of different patients (Spearman rank coefficient = 0.050; *P* = 0.6). For *msp-2*, (Spearman rank coefficient = 0.094; *P*= 0.3) and for *glurp* genes (Spearman rank coefficient = 0.036; *P* = 0.7) there was also no correlation between MOI and age group (Table 5)

**Table 5:** Distribution of *msp-1* (block-2), *msp-2* (block-3) and glurp RII allelic type among different age group of *P. falciparum* infected patients from KP, Pakistan.

| Genes | Allelic type | <5 | 5-20 | 21-40 | 41-60 | >60 | Total |
| --- | --- | --- | --- | --- | --- | --- | --- |
| msp-1 | K1 | 01(1.3) | 0 | 07 (9.3) | 08 (10.7) | 0 | 16 (21.3) |
|  | MAD20 | 01(1.3) | 04 (5.3) | 01(1.3) | 00 | 0 | 06 (8.00) |
|  | RO33 | 02(2.7) | 01 (1.3) | 03 (4.0) | 01 (1.3) | 01 (1.3) | 08 (10.7) |
|  | K1/MAD20 | 0 | 03 (4.0) | 02 (2.7) | 04 (5.3) | 0 | 09 (12.0) |
|  | K1/RO33 | 01(1.3) | 01(1.3) | 05 (6.7) | 03 (4.0) | 0 | 10 (13.3) |
|  | MAD20/ RO33 | 01(1.3) | 01(1.3) | 06 (8.0) | 03 (4.0) | 0 | 11 (14.7) |
|  | K1/MAD20/RO33 | 0 | 02(2.7) | 08 (10.7) | 05 (6.7) | 0 | 15 (20.0) |
|  | Total | 06 (8.0) | 12 (16.0) | 32 (42.7) | 24 (32.0) | 01 (1.3) | 75 (100) |
|  | Total K1 | 02 (33.3) | 06 (50.0) | 22 (68.7) | 20 (83.3) | 0 | 50 (66.7) |
|  | Total MAD20 | 02 ((33.3) | 10 (83.3) | 17 (53.1) | 12 (50.0) | 0 | 41 (54.7) |
|  | Total RO33 | 04 (66.7) | 05 (41.7) | 23 (71.9) | 12 (50.0) | 01 | 44 (58.7) |
|  | Multiclonal Isolates | 02 (2.7) | 07 (9.3) | 22 (29.3) | 15 (20.0) | 0 | 45 (60.0) |
|  | MOI | **1.4** | **1.7** | **1.9** | **1.9** | **1.0** | **1.8** |
| msp-2 | FC27 | 01(1.5) | 04 (6.0) | 08 (12.0) | 08 (12.0) | 01 (1.5) | 22 (32.8) |
|  | 3D7 | 01 (1.5) | 03 (4.5) | 11 (16.4) | 05 (7.5) | 0 | 20 (29.9) |
|  | FC27/3D7 | 01 (1.5) | 04 (6.0) | 09 (13.4) | 11 (16.4) | 0 | 25 (37.3) |
|  | Total | 03 (4.5) | 11 (16.4) | 28 (41.8) | 24 (35.9) | 01(1.5) | 67 (100) |
|  | Total FC27 | 02(66.7) | 08 (72.7) | 17 (60.7) | 19 (79.2) | 01 (100) | 47 (70.1) |
|  | Total 3D7 | 02(66.7) | 07 (63.6) | 20 (71.4) | 16 (66.7) | 0 | 45 (67.2) |
|  | Multiclonal Isolates | 01(1.5) | 04 (6.0) | 09 (13.4) | 11 (16.4) | 0 | 25 (37.3) |
|  | MOI | **1.4** | **1.4** | **1.3** | **1.5** | **1.0** | **1.4** |
|  | Overall MOI | **1.4** | **1.6** | **1.6** | **1.7** | **1.0** | **2.6** |

| Glurp | 07 | 10 | 21 | 23 | 01 | 62 |
| --- | --- | --- | --- | --- | --- | --- |
| MOI | **1.1** | **1.0** | **1.05** | **1.0** | **1.0** | **1.03** |

The distribution of *msp-1, glurp* genes and their sub-allelic families according to parasitaemia having no significant correlation differences observed. (Spearman rank 0.205; *P* = 0.060) (0.026; *P* = 0.814) respectively while MOI and parasite density for *msp-2* was significantly correlated (Spearman rank coefficient 0.250; *P* = 0.021). The MOI and parasite density of all the markers collectively (*msp-1, msp-2* and *glurp*) was not significantly correlated (*P* = 0.23). For *msp-1* lower MOI observed in the range of 5001-10000 parasite density, while higher MOI observed in the range of 50-5000 parasitaemia. For *msp-2* lower MOI detected for 50-5000 parasitaemia while higher MOI detected for more than 10000 parasitaemia. The distribution of different alleles of *msp-1, msp-2* and *glurp* according to parasitaemia is shown in (Table 6)

**Table 6:** Distributions of *msp-*1 (blocks-2), *msp-2* (block-3) and glurp RII allelic types among parasitaemia groups in the study area.

| **msp-1** | **50 – 5000** | **5001- 10000** | **>10000** | **Total** |
| --- | --- | --- | --- | --- |
| **K1** | 0 | 10(13.3) | 6(8.0) | 16(21.3) |
| **MAD20** | 1(1.4) | 1(1.4) | 4(5.3) | 06(8.00) |
| **RO33** | 0 | 5(6.7) | 3(4.0) | 08(10.7) |
| **K1/MAD20** | 3(4.0) | 2(2.6) | 4(5.3) | 09(12.0) |
| **K1/RO33** | 1(1.4) | 6(8.0) | 3(4.0) | 10(13.3) |
| **MAD20/ RO33** | 1(1.4) | 2(2.6) | 8 (10.7) | 11(14.7) |
| **K1/MAD20/RO33** | 2(2.6) | 2(2.6) | 11(14.7) | 15(20.0) |
| **Total** | 8(10.8) | 28(37.2) | 39(52.0) | 75(100) |
| **Total K1** | 6 (75.0) | 20(71.4) | 24(61.5) | 50(66.7) |
| **Total MAD20** | 7 (87.5) | 7(25.0) | 27(69.2) | 41(54.7) |
| **Total RO33** | 4 (50.0) | 15(53.6) | 25(64.1) | 44(58.7) |
| **Muticlonal isolates** | 07(9.3) | 12(16.0) | 26(34.6) | 45(60.0) |
| **MOI** | **2.1** | **1.5** | **1.7** | **1.8** |
| **msp-2** |  |  |  |  |
| **3D7** | 3(4.5) | 8(11.9) | 11(16.4) | 22 (32.8) |
| **FC27** | 2 (3.0) | 10 (15.0) | 08 (11.9) | 20 (29.9) |
| **3D7+FC27** | 2 (2.9) | 7(10.4) | 16(23.9) | 25 (37.3) |
| **Total** | 7 (8.9) | 25(37.3) | 35 (53.7) | 67(100) |
| **Total 3D7** | 5 (66.7) | 15(60.0) | 27(75) | 47 (70.1) |
| **Total FC27** | 4 (50.0) | 17(68.0) | 24 (69.4) | 45 (67.2) |
| **Multiclonal isolates** | 02(1.5) | 07(10.4) | 16(23.9) | 25 (37.3) |
| **MOI** | **1.2** | **1.3** | **1.5** | **1.4** |
| **Overall MOI** | **1.6** | **1.4** | **1.6** | **2.6** |
| **glurp** | 9(15) | 20(33.4) | 31(51.6) | 60 (100) |
| **MOI** | **1.0** | **1.0** | **1.0** | **1.03** |

**Table 7:**
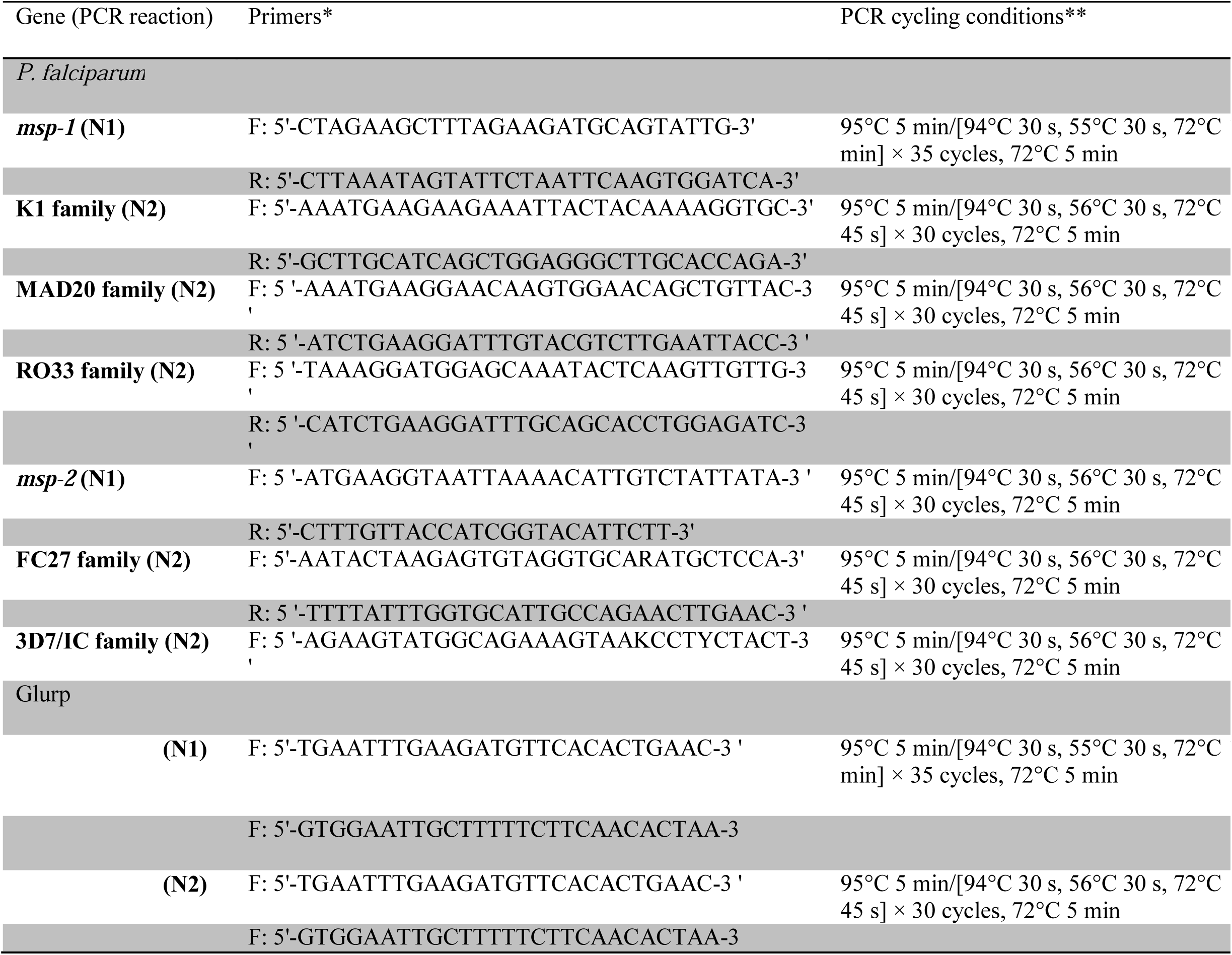
Different sequences of the primers used to amplify *msp1, msp2* and *glurp* genes of *P. falciparum* isolates from KPK, Pakistan.

## Discussion

The present study provides the most detailed analysis and assessment of genetic diversity and polymorphism of *P*.*falciparum* in Khyber Pakhtunkhwa, Pakistan. The purpose of this study was to determine the genetic diversity and population structure of *Plasmodium* species using different polymorphic regions of *msp-1, msp-2, and glurp* in ninedifferent districts of our study site. Genetic diversity and polymorphism play an important role in the acquisition of anti-malaria parasite immunity (Chenet *et al*., 2008 & Zekari *et al*., 2005) therefore to identify those genotypes which circulating in different geographical locations and areas would facilitate the development of effective control strategies.

In our study we analyzed, genotyping of *msp-1, msp-2* and *glurp* genes. The diversity of *msp1, msp2* alleles were higher than *glurp*. This result is the line with previous reports (Congpuong *et al*., 2014 & Soe *et al*., 2017). Among the allelic families of *msp1* genes, total of 32 allelic types are detected in which K1 was the predominant allelic type, similar to reports from Southwest Ethiopia (Mohammad*et al*., 2015), Cote d’ Ivoire and Gabon (Yavo *et al*., 2016) Central Africa, Gabon, Benin and Ghana (Wezam, 1994, Snounou *et al*., 2002 & Gosi *et al*., 2013) where K1 was the highly prevalent allelic family. In contrast the isolates from Indonesia (Sorontou *et al*., 2015), Malaysia (Mohd *et al*., 2016) and Sudan (Hamid *et al*., 2016) where MAD20 allele of *msp-1* were prevalent. So the variation in the prevalence of *msp-1* alleles between different studies likely reflects the variation in geographical locations (Yavo *et al*., 2016 & Farnert*et al*., 2008).

For *msp-2*, the total of 30 alleles types were detected, the alleles belonging to 3D7 family were predominant. This data is same as reported from Kenya (Takala *et al*., 2002), Congo Brazzaville (Mayengue *et al*., 2011), Peru (Chenet *et al*., 2008) Iran (Zakeri *et al*., 2005) and other sub-Saharan African countries (Mwingira *et al*., 2011) While in contrast, Osogbo Nigeria (Ojurongbe *et al*., 2011) and North-Eastern Myanmar (Yuan *et al*., 2013) the situation is opposite. For *glurp* gene, the RII region of the *glurp* was also show high degree of polymorphism in the parasite population with nine different allelic fragments size detected. The high level of *glurp* alleles shows a high malaria endemicity which is also in the agreement with the previous data (Mwingira *et al*., 2011 & Kumar*et al*., 2014).

For *msp-1, msp-2* and *glurp*the high level of genetic diversity is compatible with high level of malaria transmission. A total of 30, 32 and 9 distinct alleles for *msp-1* and *msp-2* and *glurp* respectively were obtained from the isolates. In our study the genetic diversity is higher than that in the other countries like Nigeria (*msp-1*: 5; *msp-2*: 15), (Olasehinde *et al*., 2012), Brazzaville of the Republic of Congo (*msp-1*: 15; *msp-2*: 20) (Gueye *et al*., 2018)and Bangui of Central African Republic (*msp-1*: 17; *msp-2*: 25) (Dolmazon *et al*., 2008) but lower than Gabon (*msp-1*: 39; *msp-2*: 27) (Ngomo *et al*., 2018). For *glurp* genes the same result shows by Northwest Ethiopia (9 genotypes) (Mohammed*et al*., 2018) while in India (8 genotypes) were recorded (Gupta*et al*., 2014).In the agreement with previous reports, the RII region of *glurp* gene was also polymorphic with (11 genotypes) in Southwestern Nigeria (Funwei *et al*., 2008) and (14 genotypes) in Sub-Sahara Africa (Mwingira *et al*., 2011) which shows the higher number *P. falciparum* genotypes than our study.

The overall mean multiplicity of infection was 2.6, when considering of *msp-1, msp-2* and *glurp* genes separately, the multiplicity of infection was (MOI= 1.8) for *msp-1*, (MOI=1.4) for *msp-2*and (MOI= 1.03) for *glurp* gene. MOI reported in this study was higher than other countries like Malaysia where the MOI of *P. falciparum* infection for *msp-1* was (MOI = 1.37) and for *msp-2* (MOI = 1.20) (Atroosh *et al*., 2011). But in our study the value of MOI for *msp-2* was lower in Cote d’ Ivoire (MOI = 2.8) (Silue *et al*., 2006). The overall MOI in our study was (2.6), this result was similar with Northwest Ethiopia where the mean MOI was (2.6) (Muhammad *et al*., 2018).Thisresults value is higher than that reported in other countries, including Southern Ghana (1.17-1.48)

Adjah *et al*., 2018 reported the Republic of Congo (1.7) (NdongNgomo *et al*., 2018), Southwest Ethiopia (1.8) (Mohammed *et al*., 2015) and Republic of Congo (2.2) (Mayengue *et al*., 2011) While in Guinea the value of overall MOI was (5.51) (Chen *et al*., 2018), Southwestern Nigeria (2.8) (Funwei *et al*., 2018) and in Mauritania (3.2) (Salam *et al*., 2014) which is higher than our values.This discrepancy may be due to change in geographical areas their transmission pattern and also samples population determination. When higher the malaria transmission level there will be greater chance to get a higher MOI and mean number of alleles per locus.

In our study we found (81%) of samplesharboredmulticlonal infection (having more than one allele type) which is in the line with previous studies from Mauritania which is (82.3%) (Salem *et al*., 2014), Brazzaville Republic of Congo (83%) (Mayengue *et al*., 2011) and Iran which is (87%) (Zakeri *et al*., 2005) while in contrast (70%) from Nigeria (Funwei *et al*., 2018) (59%) from Southwest Ethiopia (Mohammad *et al*., 2015) and (62%) from Sudan (Hamid *et al*., 2013) found a lower frequency of samples with multiclonal infection. High level prevalence of multiclonal infection of *msp-1* and *msp-2* genotypes was observed in our study. This high level of genetic diversity is an indication that the parasite population size remained high enough to allow better mixing of different genotypes and also human migration bring to change in genetic diversity by introducing additional number of parasites genotype (Mita *et al*., 2014)

Age is a key factor to affect MOI in *P. falciparum* infection and it also involve in the acquisition of immunity against *P. falciparum* species (Auburn *et al*., 2017). In young patients low level of immunity may be the major factor contributing to their vulnerability to control the infection (Vardo-Zalik *et al*., 2013 & Yuan*et al*., 2013). The high level of malarial transmission in the region may be expected to lead to a higher risk of severe malaria in younger patients where immunity is lower (Escalante *et al*., 2015)

In our study MOI was not correlated by patient age as shown similar results in other countries like Sudan (Bakhi *et al*., 2015), Ethiopia (Wezam, 1994), Benin (Ogouyèmi-Hounto *et al*., 2013) and Senegal (Vafa *et al*., 2008) which suggests that the MOI is not directly related to the period of acquisition of immunity in asymptomatic patients but reflects the exposure of subjects to malaria in the endemic area but having a contrast results to Central Sudan (Hamid MM *et al*.,2013), Burkina Faso (Soulama *et al*., 2009) and Bioko Island, Equatorial Guinea (Chen *et al*., 2018).

A positive correlation between MOI and parasite density for *msp-2* was analyzed in this study including Congo Brazzaville (Mayengue *et al*., 2011), (Vafa *et al*., 2008) and West Uganda (Peyerl-Hofmann *et al*., 2010) (*P* = 0.01) which show similar results, while contrast study show by South Benin (Ogouyemi-Hounto *et al*., 2013).

When the parasite density was categorized into three groups (50 -5000) (5001 - 10000) (>10000 parasite/µl) we found that there was not significantly influenced by MOI (multiplicity of infection) (*P* = 0.23). Different studies have shown a relationship between the MOI and parasite density but in our results the MOI did not increase accordingly to increase with parasite densities. Our results were similar in Benin (Ogouyèmi-Hounto *et al*., 2013) and Nigeria (Ojurongbe *et al*., 2011) while contrast result show by Congo, Brazzaville (Mayengue *et al*., 2011) and West Uganda (Peyerl-Hoffmann *et al*., 2001)

These results are consistent with many reports demonstrating that high parasite densities increase the probability of detecting concurrent clones in individuals (Peyerl-Hoffmann *et al*., 2010 & Färnert *et al*., 2001). This finding shows that high malaria transmission and also parasite density may have a strong association with the genetic diversity of *P. falciparum* in Khyber Pakhtunkhwa province of Pakistan.

## Conclusion

The present study shows that field isolates of nine different districts of Khyber Pakhtunkhwa are highly diverse in respect of *msp-1* (block-2), *msp-2* (block-3) and glutamate rich protein.This study provides basic information on the genetic diversity and multiple infection of *P. falciparum* (*msp1, msp2* and *glurp*) isolates from study site of Pakistan. This information will be baseline data for future studies on dynamics of parasite transmission and for the purpose to evaluate the malaria control interventions in our study area of Pakistan.

